# Non-CG DNA methylation modulates hypocotyl elongation during thermormorphogenesis

**DOI:** 10.1101/2024.09.25.614994

**Authors:** Maián Garro, Eleonora Greco, Gustavo J. Vannay, Aleksandra Leonova, Leonardo Bruno, Matías Capella

## Abstract

Plants adapt to warm environments through physiological and morphological changes termed thermomorphogenesis, which involve transcriptional reprogramming exerted mainly by PHYTOCHROME INTERACTING FACTOR 4 (PIF4). Fluctuating temperatures can also influence the patterns of cytosine DNA methylation, thereby influencing gene expression. However, whether these epigenetic changes provide an adaptative advantage remains unclear. Here, we provide evidence that DNA methylation is required to regulate thermomorphogenesis. Hypomethylated *drm1 drm2 cmt3* mutants or seedlings treated with 5-azacytidine to block DNA methylation exhibit reduced hypocotyl growth at warm temperatures, primarily due to impaired cell elongation. Moreover, DNA hypomethylation compromises auxin biosynthesis and transport in response to warmth, partially by reducing PIF4 protein levels. Notably, the loss of DNA methylation leads to increased expression of *SUPPRESSOR OF drm1 drm2 cmt3* (*SDC*), which in turn restricts hypocotyl elongation during thermomorphogenesis. Finally, we demonstrate that DNAme regulates the inhibition of *SDC* expression to promote gibberellin biosynthesis. Our findings underscore the critical role of DNA methylation in modulating gene expression in response to temperature fluctuations and provide new insights into the epigenetic regulation of thermomorphogenesis.

**Highlights:** DNA methylation regulates the expression of key genes involved in auxin and gibberellin metabolism, to ensure hypocotyl growth in response to warm temperatures.

## Introduction

Elevated temperatures are a main environmental factor affecting plant growth and crop productivity. Plants evolved sophisticated mechanisms to sense, adapt and ultimately survive in environments with fluctuating temperatures. At warm temperatures, plants undergo a series of physiological and morphological adaptations, collectively known as thermomorphogenesis, to mitigate exposure to potentially harmful conditions (Casal and Balasubramanian, 2019). These adaptations include hypocotyl and petiole elongation, increased leaf hyponasty, and accelerated flowering. The bHLH-type transcription factor PHYTOCHROME INTERACTING FACTOR 4 (PIF4) is a key regulator of the transcriptional reprogramming that results in the developmental response induced by thermomorphogenesis in Arabidopsis (Jin *et al*., 2020).

Interestingly, warming influences *PIF4* expression, protein levels, and its function as a transcription factor (Delker *et al*., 2022). For instance, ELONGATED HYPOCOTYL 5 (HY5) represses *PIF4* expression and competes for a place in the promoters of PIF4-targeted genes upon warming (Delker *et al*., 2014; Toledo-Ortiz *et al*., 2014). Additionally, various phytohormones play a critical role in regulating PIF4-mediated thermomorphogenesis. Specifically, PIF4 directly activates the expression of phytohormone biosynthesis genes, such as *YUCCA8*, leading to increased auxin levels in aerial tissues and promoting hypocotyl elongation (Sun *et al*., 2012). Moreover, gibberellins (GA) facilitate the degradation of DELLA proteins, which would otherwise bind to PIF4 and either inhibit its DNA-binding capacity or reduce its protein stability, thereby limiting hypocotyl elongation during thermomorphogenesis (De Lucas *et al*., 2008; Li *et al*., 2016; Park *et al*., 2020).

In addition to the transcriptional control influenced by PIF4, gene expression during thermomorphogenesis can be modulated by epigenetic mechanisms, which alter the accessibility of transcriptional machinery without changing DNA sequences. One key epigenetic modification is the addition of a methyl group to cytosine at the 5’ position, known as DNA methylation (DNAme). In plants, DNAme occurs at cytosine bases in CG, CHG, and CHH contexts (where H is A, T, or C) and plays a crucial role in maintaining genome stability and regulating gene expression. DNAme is established by the RNA-directed DNA methylation (RdDM) pathway, which requires the methyltransferase DOMAINS REARRANGED METHYLTRANSFERASE 2 (DRM2) and proteins involved in the generation of small interfering RNAs (siRNAs), such as DICER-LIKE 3 (DCL3) (Matzke and Mosher, 2014). Maintenance of DNA methylation depends on the sequence context. The methyltransferase MET1 and VARIANT IN METHYLATION (VIM) proteins maintain CG methylation, while CHG methylation is preserved by CMT3 and, to a lesser extent, by CMT2. Finally, the maintenance of CHH methylation requires the RdDM pathway or CMT2, depending on the sequence environment. The plant SU(VAR)3-9 homologs SUVH4 to SUVH6 are methyltransferase enzymes deposit a methyl group on histone H3 at Lys9 (H3K9me). These proteins are recruited to DNA by CHG methylation, and reinforces DNAme by recruiting the DNA methyltransferases CMT2 and CMT3 (Xu and Jiang, 2020). Finally, DNAme can be passively lost during DNA replication, or actively removed by DNA glycosylases, such as REPRESSOR OF SILENCING (ROS1), TRANSCRIPTIONAL ACTIVATOR DEMETER (DME), DEMETER-LIKE PROTEIN 2 (DML2), and DML3 (Zhang *et al*., 2022).

In plants, DNAme primarily targets transposons and other repetitive DNA elements across all contexts, effectively silencing their expression (Zhang *et al*., 2006; Lister *et al*., 2008). When a transposon or a repetitive sequence is situated near or within a gene, DNAme can activate or repress gene expression, thereby influencing plant development and adaptation to diverse environmental challenges (Kinoshita *et al*., 2007; Stuart *et al*., 2016; Roquis *et al*., 2021; Arce *et al*., 2023). For example, CG DNAme represses the auxin biosynthesis *YUCCA2* gene expression, and its removal promotes *YUCCA2* induction during thermomorphogenesis (Fonouni-Farde *et al*., 2022; Forgione *et al*., 2022). Moreover, CHG and CHH DNAme occurring at tandem repeats within the promoter of the F-box-containing gene *SUPPRESSOR OF drm1 drm2 cmt3* (*SDC*) silences its expression, preventing the development of curled leaves and reduced growth (Henderson and Jacobsen, 2008). The curly-shaped leaves exhibited by plants deficient in non-CG DNAme are also partially attributed to the misregulation of genes critical for auxin homeostasis (Forgione *et al*., 2019). Moreover, the absence of DNAme leads to the repression of genes involved in leaf shape development, such as *KNAT6*, by promoting the deposition of the silencing epigenetic mark H3K27me3 (Forgione *et al*., 2022). Despite its crucial functions in regulating plant development and responses to environmental conditions (Zhang *et al*., 2018; Liu and He, 2020), the potential role of DNAme in the CHG and CHH contexts in modulating thermomorphogenesis remains unexplored.

Here, we report that non-CG DNAme is required to regulate plant response to warm temperatures. We show that *drm1 drm2 cmt3* mutants and seedlings treated with 5-azacytidine to induce DNA hypomethylation exhibit reduced hypocotyl elongation during thermomorphogenesis due to impaired cell elongation. Additionally, our data reveal that DNAme positively influences auxin levels at the hypocotyl by promoting both auxin biosynthesis and transport. Furthermore, DNAme helps maintain PIF4 protein stability, likely by inducing GA biosynthesis. Importantly, SDC mediates the effects of DNA hypomethylation on hypocotyl elongation at warm temperatures. We propose that DNAme represses *SDC* expression to ensure auxin and GA homeostasis, facilitating plant adaptation to elevated environmental temperatures.

## Materials and Methods

### Plant material

Wild type and other lines of plants used in this study were in Arabidopsis thaliana L. Heynh. ecotype Columbia (Col-0) or Landsberg *erecta* (L*er*). The mutant seeds *drm1-2 drm2-2 cmt3-11* (*ddc*; CS16384), *dcl2-1 dcl3-1 dcl4-2t* (*dcl234*; CS16391), *suppressor of drm1 drm2 cmt3* (*sdc*; SALK_017593), *drm1-2 drm2-2 kyp-6* (CS16388), and *pif4-2* (CS66043) in the Col-0 ecotype background, and the quintuple *della* mutant (*gai-t6 rga-t2 rgl1-1 rgl3-1 rgl2-1*), and seedlings overexpressing either *RGA* (*35S::TAP-RGA*Δ*17*; CS16292) or *GAI* (*35S::TAP-GAI*Δ*17*; CS16294) in the L*er* ecotype were obtained from the Arabidopsis Biological Resource Center (ABRC; Columbus, OH, USA; http://www.arabidopsis.org). The mutant seeds *met1-3* (Saze *et al*., 2003) were generously provided by Prof. Dr. Frédéric Berger from the Gregor Mendel Institute of Molecular Plant Biology, Vienna, Austria. The *drm1-2 drm2-2 cmt2-3 cmt3-11* (*ddc2c3*) and *CsVMV::SDC-HA* seeds (Tian *et al*., 2021) were kindly supplied by Prof. Lei Wang, Institute of Botany, The Chinese Academy of Sciences, Beijing, China. The mutant seeds *ros1-4 dml2-2 dml3-2* (*rdd-2*), *dme^DD7pro^*, and *ros1-4 dml2-2 dml3-2 dme^DD7pro^* (*drdd*) (Zeng *et al*., 2021) were kindly provided by Dr. Daisuke Miki from the Shanghai Center for Plant Stress Biology, Shanghai, China. The *pPIF4::PIF4-GFP pif4-101* seeds (Pucciariello *et al*., 2018) were generously supplied by Dr. Jorge J. Casal from Instituto de Investigaciones Fisiológicas y Ecológicas Vinculadas a la Agricultura (IFEVA), Buenos Aires, Argentina. The seeds of *A. thaliana* ecotype Sha (CS76382) and No-0 (CS77128) were kindly provided by Dr. Pablo A. Manavella from the Instituto de Agrobiotecnología del Litoral, Santa Fe, Argentina.

### Plant growth conditions and treatments

*Arabidopsis thaliana* plants were grown on soil in a 21-23 °C growth chamber under a long-day photoperiod (16 h light, 110 μE m^−2^ s^−1^/8 h dark) in 8 x 7 cm. Seeds for hypocotyl length evaluation were surface-sterilized by treatments with 70 % EtOH and 10 % sodium hypochlorite, washed, and stratified at 4 °C for 3 days to obtain homogeneous germination. Unless stated otherwise, seedlings were grown in Petri dishes containing half-strength Murashige and Skoog (MS/2) basal medium (Murashige and Skoog, 1962) supplemented with vitamins (PhytoTechnology Laboratories) and 0.9 % agar, at 22°C under long-day conditions (16 h light, 110 μE m^−2^ s^−1^/8 h dark) for 3 days, and transferred to 29 °C or maintained at 22 °C for 3 additional days. For *Nicotiana benthamiana*, surface-sterilized seeds were grown in Petri dishes containing MS/2 supplemented with vitamins and 1 % agar under long-day conditions for 7 days, and transferred to 29 °C or maintained at 22 °C for 4 additional days. To measure petiole length, plants were grown on soil at 22°C for 9 days, and transferred to 29 °C or maintained at 22 °C for 6 additional days. For treatments, seeds were grown in MS/2 medium containing 75 µM 5-azacytidine (Goldbio, St. Louis, MO, USA), 0.1 or 0.5 µM picloram (Duchefa Biochemie, Haarlem, the Netherlands), 10 µM GA_3_ (Duchefa Biochemie, Haarlem, the Netherlands) or 10 nM PAC (Sigma-Aldrich, St. Louis, MO, USA) at 22°C under long-day conditions for 3 days, and transferred to 29 °C or maintained at 22 °C for 3 additional days. The 5-azacytidine, picloram, GA_3_, and PAC stock solutions were prepared according to the manufacturer’s recommendation. Note that treatments at 29°C were performed using ARALAB Fitoclima 600, LED light.

### Measurement of hypocotyl, cell, and petiole length

As previously reported (Capella *et al*., 2015), hypocotyl length was determined by measuring the distance from the most basal root hair to the ‘V’ shape made by the cotyledons. Cell length was determined by observing seedlings under a confocal microscope. For petiole length, the fourth leaves of 16-day-old plants from each genotype were dissected from the rosette, scanned, and analyzed. Hypocotyl length, cell length, and petiole length were determined using Fiji/ImageJ software (Schindelin *et al*., 2012).

### Histochemical GUS staining

*In situ* GUS staining was performed as previously described (Jefferson *et al*., 1987). Seedlings were grown on MS/2 basal medium with or without 75 µM 5-azacytidine at 22 °C under long-day conditions for 3 days and transferred to 29 °C or maintained at 22 °C for 3 extra days. Seedlings were immersed in GUS staining buffer (1 mM 5-bromo-4-chloro-3-indolyl-b-glucuronic acid solution in 100 mM sodium phosphate, pH 7, 0.1 % (v/v) Triton X-100, 100 mM potassium ferrocyanide, and 100 mM potassium ferricyanide), vacuum was applied for 5 min, and then plants were incubated at 37 °C overnight. Chlorophyll was cleared from green plant tissues by immersing them in 100 % ethanol.

### Fluorescence microscopy

For confocal imaging, *pPIF4::PIF4-GFP* and *DR5::GFP* seedlings were grown on MS/2 basal medium with or without 75 µM 5-azacytidine at 22 °C under long-day conditions for 3 days and transferred to 29 °C or maintained at 22 °C for 3 additional days. Seedlings were examined and imaged with a confocal inverted microscope (Confocal LEICA TCS SP8), using a 10× (for *DR5::GFP*) or a 20× (for *pPIF4::PIF4-GFP*) objective, a 514 nm excitation line laser for GFP (50% intensity), and appropriate emission at 498 nm − 532 nm bandpass filters. All images were captured using identical microscope settings. Fluorescence intensity was measured from maximum Z-projections within defined areas of individual nuclei in *pPIF4::PIF4-GFP* seedlings grown at 29°C, with or without 5-azacytidine. A minimum of 10 nuclei from 4 independent biological samples were analyzed per condition Subsequent processing and analyses of the images were performed in Fiji/ ImageJ (Schindelin *et al*., 2012).

### RNA isolation and analyses

Transcript levels were evaluated by RT-qPCR. Samples were prepared from seedlings grown on MS/2 basal medium at 22 °C under long-day conditions for 3 days and transferred to 29 °C or maintained at 22 °C for 3 additional days. At least 80 seedlings from each genotype were harvested at ZT2 and processed for RNA extraction, and 3-4 biological replicates were used for each RT-qPCR experiment. The total RNA was purified from seedlings using Trizol^®^ reagent (Invitrogen) according to the manufacturer’s instructions. One μg of RNA was reverse-transcribed using oligo(dT)_18_ and M-MLV reverse transcriptase II (Promega). Quantitative real-time PCR (qPCR) assays were performed using a StepOne equipment (Applied Biosystems); each reaction contained a final volume of 20 μl that included 2 μl of SyBr green (4×), 8 pmol of each primer, 2 mM MgCl_2_, 10 μl of a 1/20 dilution of the RT reaction and 0.1 μl of Taq Polymerase (Invitrogen), using standard protocols (40 to 45 cycles, 60 °C annealing). Fluorescence was quantified at 72 °C. Specific primers for each gene were designed and are listed in Supplementary Table S1. The expression levels were normalized using *PP2A.A3* (AT1G13320) or *SAND* (AT2G28390) as reference genes, and quantification was carried out using the ΔΔCt method (Pfaffl, 2001).

### RNA-seq analysis

We reanalyzed public RNA-Seq data for WT seedlings grown at 22°C and 29°C previously reported (He *et al*., 2022). RNA-seq reads were analyzed on the Galaxy platform (Abueg *et al*., 2024). Briefly, adapter sequences were removed from the reads using Trimmomatic (Galaxy Version 0.39+galaxy2). The quality-filtered reads were aligned to the *Arabidopsis thaliana* reference genome (TAIR10) using RNA STAR (Galaxy Version 2.7.11a+galaxy1), employing a maximum intron length of 20,000 bp and guided by the gene and exon annotation from Araport V11. The read counts on each gene were then calculated using featureCounts (Galaxy Version 2.0.3+galaxy2). Differential expression analysis was performed using DESeq2 (Galaxy Version 2.11.40.8+galaxy0). Bigwig coverage files were generated using bamCoverage (Galaxy Version 3.5.4+galaxy0), and subsequently used for plotting with pyGenomeTracks (Galaxy Version 3.8+galaxy2). RNA-seq data was previously deposited in the Gene Expression Omnibus (GEO) database under the accession number GSE181292.

### Statistics and reproducibility

Representative results of at least two independent experiments were presented in all of the figure panels. Analyses of the variance were performed, and pairwise differences were evaluated with Tukey’s *post hoc* test using R statistical language (R Development Core Team, 2008); different groups are marked with letters at the 0.05 significance level. For all error bars, data are mean ± S.E.M. *P* values were generated using two-tailed Student’s *t*-tests; N/S, *P* ≥ 0.05, *\*P* ≤ 0.05, *\*\*P* ≤ 0.01, *\*\*\*P* ≤ 0.001.

### Accession numbers

CMT2 (AT4G19020); CMT3 (AT1G69770); DRM1 (AT5G15380); DRM2 (AT5G14620); MET1 (AT5G49160); PIF4 (AT2G43010); SDC (AT2G17690)

## Results

### DNA methylation is required for hypocotyl response to warm temperatures

CHG and CHH methylation maintains auxin homeostasis at 22°C, and demethylation of CG sites promotes auxin synthesis during thermomorphogenesis (Forgione *et al*., 2019; Fonouni-Farde *et al*., 2022). We thus postulated that DNAme maintenance by DRM1, DRM2, and CMT3 might contribute to plant responses to warm conditions. To test whether non-CG DNAme influences thermomorphogenesis, we measured hypocotyl length in wild-type (WT) seedlings and non-CG hypomethylated triple mutants *drm1 drm2 cmt3* (referred to as *ddc*) grown at warm temperatures conditions (see ‘Materials and Methods’). We found that the *ddc* mutants exhibited shorter hypocotyls than WT seedlings when exposed to 29°C, comparable to the *pif4* negative control (Fig. 1A,B). These results were consistent in WT and *ddc* seedlings grown at 22°C or 29°C for 5 days (Supplementary Fig. S1A). We also observed that the quadruple mutant *drm1 drm2 cmt2 cmt3* (*ddc2c3*) showed impaired hypocotyl elongation at warm temperatures (Supplementary Fig. S1B, left panel).

**Fig. 1.**
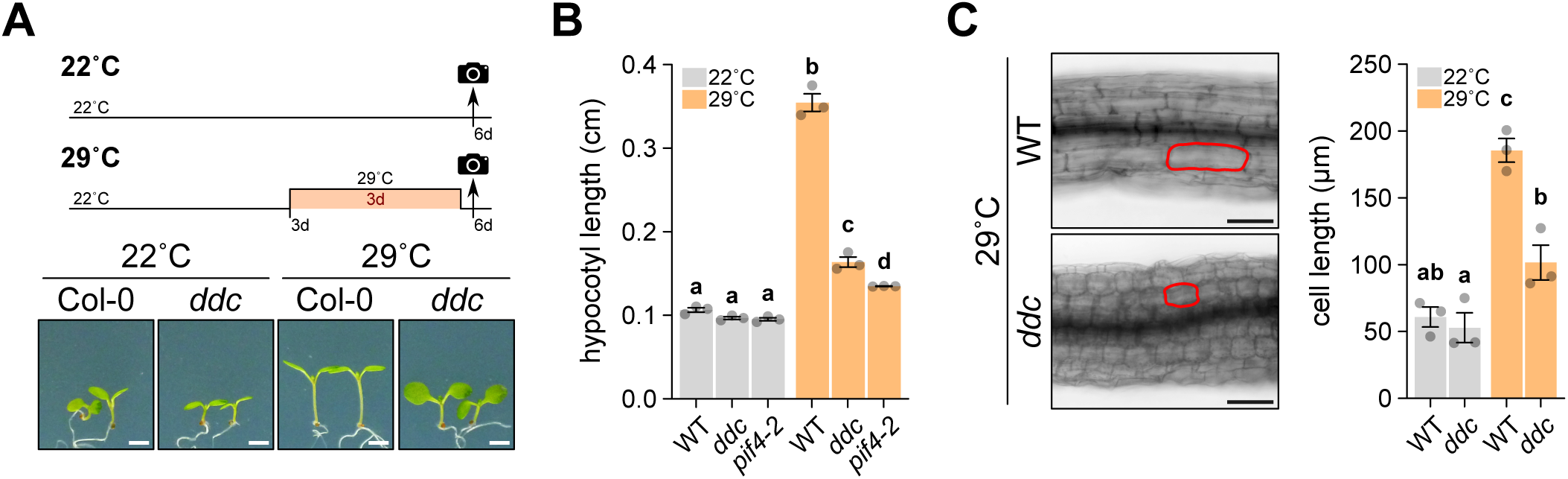
DNA methylation positively regulates warming responses in *Arabidopsis thaliana*. (A) Top, Scheme of the temperature conditions used to evaluate hypocotyl length. Bottom, illustrative photographs of WT and *ddc* seedlings grown at 22°C or 29°C. Scale bars, 2 mm. (B) Quantification of hypocotyl length of 3-day-old WT, *ddc*, and *pif4-2* seedlings transferred to 22°C or 29°C for 3 days. (C) Representative images (left) and quantification (right) of hypocotyl cell length from WT and *ddc* seedlings, grown as described in (B). Scale bars, 100 µm. For (B), and (C), data are mean ± SEM of *n* = 3 independent biological replicates. Letters denote groups with significant differences from one-way analysis of variance (ANOVA) followed by Tukey’s *post hoc* tests at *P* < 0.05.

While DRM1/2 maintains CHH methylation, CHG methylation is mostly controlled by CMT3 which requires the deposition of H3K9me2 by the methyltransferase SUVH4/KRYPTONITE (KYP) (Lindroth *et al*., 2001; Jackson *et al*., 2002). Notably, plants with a combination of loss-of-function alleles of SUVH4, DRM1, and DRM2 (*dds4*) exhibit a phenotype similar to that of the *ddc* mutants (Henderson and Jacobsen, 2008). To further confirm the role of DNAme during thermomorphogenesis, we assessed the warming response of *dds4* mutant plants. We observed that the *dds4* seedlings resembled the *ddc* mutants when grown at 29°C, showing shorter hypocotyls than WT controls (Supplementary Fig. S1B, right panel). Next, we examined whether the impaired hypocotyl elongation in the *ddc* mutants might be due to loss of DNAme at CHG (*cmt3* mutant), CHH (*drm1 drm2* mutant), or a combination of both contexts. Our findings showed that the single *cmt3* and double *drm1 drm2* mutants exhibited hypocotyl growth similar to WT controls when exposed to 29°C (Supplementary Fig. S1C). This suggests that CHG and CHH methylation act redundantly in regulating thermomorphogenesis.

DNAme is established and maintained by double-stranded RNA cleaved by dicer-like proteins into 24-nt siRNAs (Cuerda-Gil and Slotkin, 2016). In agreement with this mechanism, we observed reduced hypocotyl elongation at 29°C in the triple mutant *dcl2 dcl3 dcl4* (*dcl234*), which is impaired in 24-nt siRNA production (Henderson *et al*., 2006) (Supplementary Fig. S1D). Since DNAme is actively removed by DNA glycosylases (Penterman *et al*., 2007), we examined whether thermomorphogenesis requires the activity of these enzymes. We found that the mutants of the Arabidopsis DNA glycosylases exhibited comparable hypocotyl elongation to WT controls at warm temperatures, suggesting that active removal of DNAme is dispensable during thermomorphogenesis (Supplementary Fig. S1E). These data indicate that DNAme is required to promote the hypocotyl response to warm temperatures in Arabidopsis.

Although warm temperatures promote a moderated increase in cell division, hypocotyl growth is mainly stimulated through cell elongation mediated by elevated auxin levels (Bellstaedt *et al*., 2019; Ai *et al*., 2023). To examine whether DNAme contributes to cell elongation during thermomorphogenesis, we quantified the length of hypocotyl cells in WT and *ddc* seedlings grown at warm temperatures. Using confocal imaging, we observed that WT seedlings exhibited temperature-induced cell elongation, consistent with previous reports (Bellstaedt *et al*., 2019) (Fig. 1C). However, the *ddc* mutants showed shorter hypocotyl cells than WT controls at 29°C, suggesting that DNAme is crucial for promoting cell elongation in response to warming (Fig. 1C).

Besides promoting hypocotyl growth, increased ambient temperatures induce other physiological and morphological changes, such as petiole elongation (Casal and Balasubramanian, 2019). To investigate whether DNAme affects warming responses other than hypocotyl elongation, we examined petiole elongation in WT, *ddc*, and *met1* plants. These seedlings were grown at 22°C for six days and then either maintained at 22°C or transferred to 29°C for an additional six days. While petiole length was slightly shorter in the DNAme mutants at 22°C, all genotypes exhibited similar petiole elongation when exposed to warm temperatures, except for the *pif4* mutants as previously reported (Supplementary Fig. S2) (Koini *et al*., 2009). Together, our data suggest that DNAme is crucial for elongating the hypocotyl in response to increased ambient temperature.

### The reagent 5-azacytidine impairs hypocotyl growth during thermomorphogenesis

To strengthen our results, we analyzed the hypocotyl elongation of seedlings exposed to the DNA methyltransferase inhibitor 5-azacytidine (5-aza). Once incorporated into the genome, DNA methyltransferases become covalently bound to 5-aza at hemimethylated sites, leading to DNA hypomethylation (Christman, 2002). Similar to *ddc* mutants, we observed that 5-aza decreased hypocotyl elongation of WT seedlings grown at 29°C (Fig. 2A,B). Comparable results were obtained for other Arabidopsis accessions, such as Sha and No-0 (Fig. 2C). The pathway regulated by DNAme during thermomorphogenesis seems to be conserved across different plant species, as evidenced by the failure of 5-aza-treated tobacco plants (*Nicotiana benthamiana*) to elongate their hypocotyls when exposed to warm temperatures (Supplementary Fig. S3).

**Figure 2.**
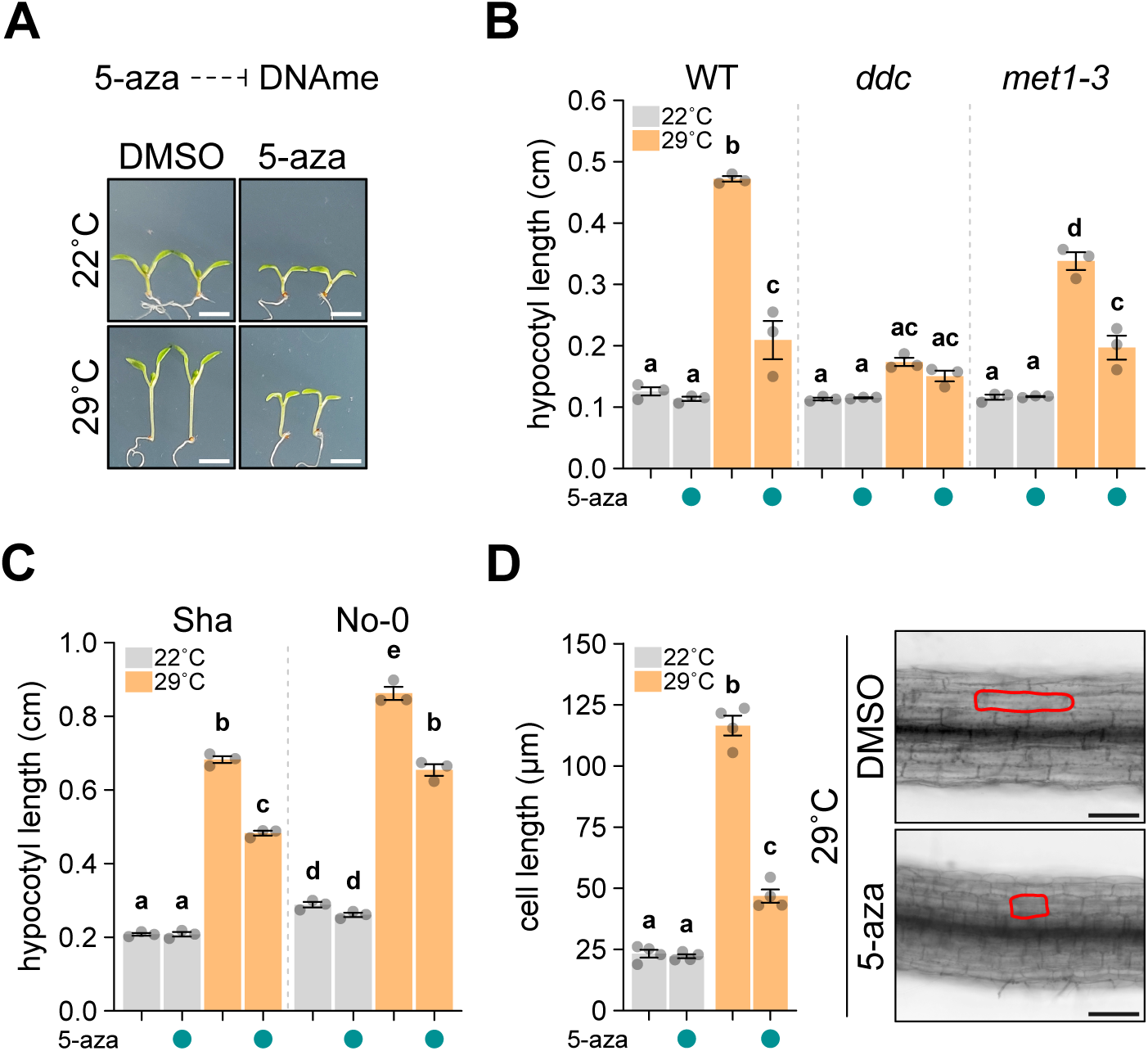
Pharmacological inhibition of DNA methylation impairs hypocotyl elongation during thermomorphogenesis. (A) Illustrative photographs of 3-day-old WT (Col-0) transferred to 22°C or 29°C for 3 additional days, in the absence or presence of 5-azacytidine 75 µM (5-aza). (B) Quantification of hypocotyl length in WT, *ddc*, and *met1-3* seedlings grown as described in (A). (C) Hypocotyl length measurements in the Arabidopsis accessions Sha and Nossen, grown as described in (A). (D) Quantification (left) and representative images (right) of hypocotyl cell length from WT seedlings grown as described in (A). Scale bars, 100 µm. For (B), (C), and (D), data are mean ± SEM of *n* = 3-4 independent biological replicates. Letters denote groups with significant differences from one-way analysis of variance (ANOVA) followed by Tukey’s *post hoc* tests at *P* < 0.05.

We then examined whether hypocotyl elongation at 29°C is also impaired in DNAme mutants treated with 5-aza. As previously reported (Fonouni-Farde *et al*., 2022), the CG methylation-deficient mutant *met1* exhibited shorter hypocotyls than WT controls, although the reduction was less severe than in *ddc* seedlings (Fig. 2B). Notably, while treatment with 5-aza further reduced hypocotyl length in *met1* mutants, the inhibition of elongation was less pronounced than in WT seedlings (Fig. 2B, right panel). In contrast, the 5-aza treatment had almost no effect on *ddc* mutants (Fig. 2A, middle panel). These results underscore the importance of non-CG DNAme for hypocotyl elongation during thermomorphogenesis.

### DNA methylation increases auxin activity in response to increased ambient temperature

The hormone auxin plays a significant role in thermomorphogenesis, with its synthesis and signaling being upregulated in response to warm temperatures. This increment in auxin activity is essential to facilitate hypocotyl elongation (Gray *et al*., 1998; Franklin *et al*., 2011; Sun *et al*., 2012). Given that the absence of DNAme impairs hypocotyl growth during thermomorphogenesis, we investigated whether this epigenetic mark impacts auxin homeostasis. To test this, we conducted histochemical analysis on seedlings bearing the ß-glucuronidase (GUS) reporter regulated by the auxin-responsive *DR5* synthetic promoter (Ulmasov *et al*., 1997), grown in the presence or absence of 5-aza at warm temperatures conditions. In agreement with previous reports (Ferrero *et al*., 2019), we observed increased GUS activity in the hypocotyls of seedlings exposed to 29°C (Fig. 3A). In contrast, treatment with 5-aza strongly inhibited GUS histochemical staining in seedlings grown at warm temperatures (Fig. 3A). Similar results were observed in *DR5::3xGFP* plants by confocal microscopy, where treatment with 5-aza reduced auxin levels in the hypocotyls of seedlings exposed to 29°C, suggesting that DNAme promotes auxin activity during thermomorphogenesis (Fig. 3B). Consistent with lower auxin levels in DNAme mutants, the addition of the synthetic auxin Picloram partially restored the hypocotyl elongation in *ddc* and *pif4* seedlings grown at 29°C (Fig. 3C, Supplementary Fig. S4). To further validate that DNAme acts upstream of auxin during thermomorphogenesis, we measured hypocotyl elongation in the auxin-overproducing mutant *yucca6-1D* after treatment with 5-aza. As previously reported (Kim *et al*., 2007), the *yucca6-1D* mutants exhibited longer hypocotyls at 22°C compared to WT controls (Fig. 3D). Notably, 5-aza-treated *yucca6-1D* mutants showed shorter hypocotyls than the untreated controls at 22°C (Fig. 3D). However, while hypocotyl elongation at 29°C relative to 22°C was lower in *yucca6-1D* than in WT, the treatment with 5-aza did not affect the *yucca6-1D* response to warm temperatures (Fig. 3E). Therefore, we conclude that DNAme modulates auxin levels to promote temperature-induced hypocotyl elongation.

**Figure 3.**
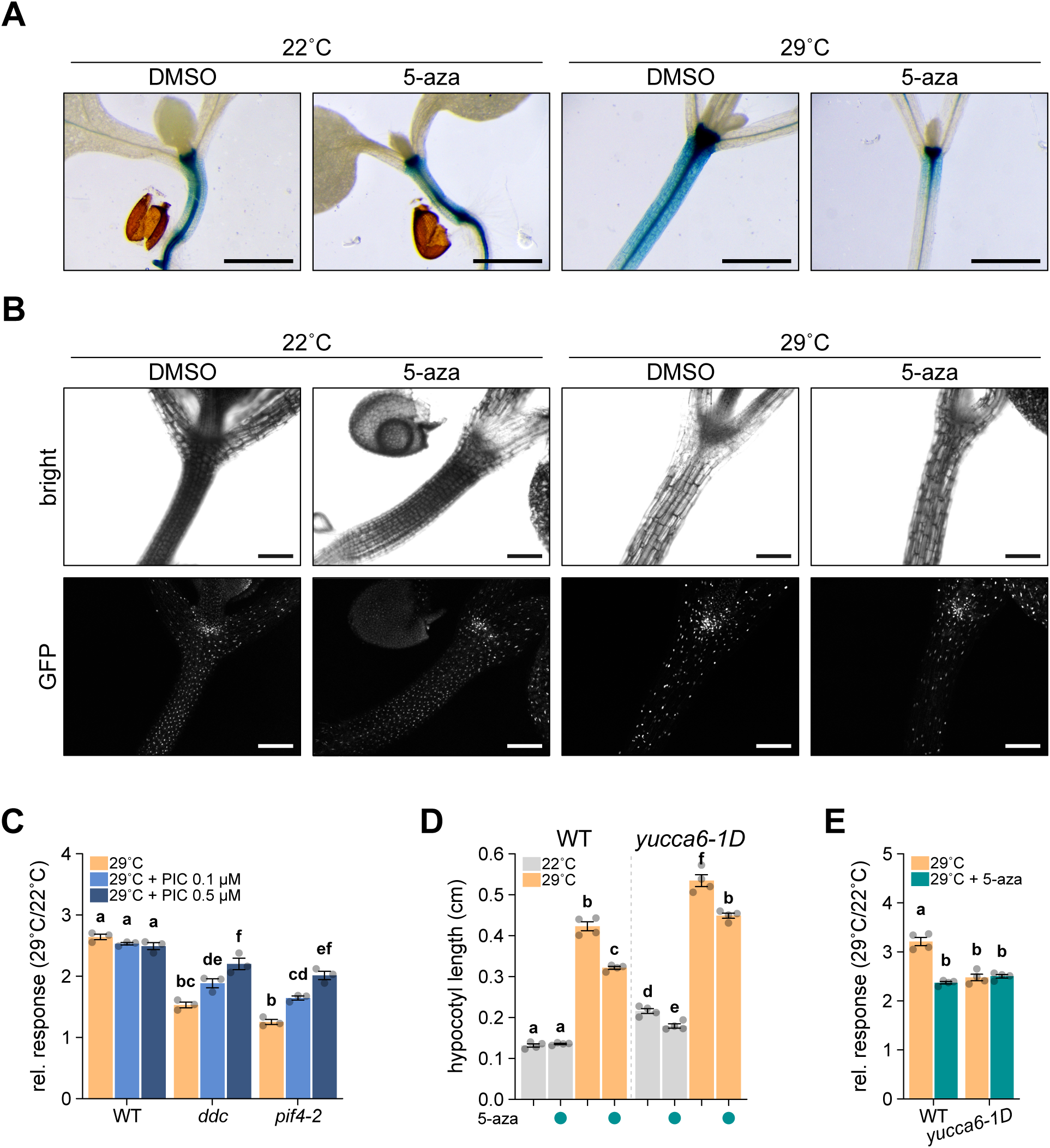
DNA methylation promotes warm-induced auxin levels. (A) Illustrative pictures from GUS histochemistry of 3-day-old *DR5::GUS* seedlings transferred to 22°C or 29°C for 3 additional days, in the absence or presence of DNA methyltransferases inhibitor 5-aza (75 µM). Scale bars, 1 mm. (B) Representative pictures of live-cell imaging of *DR5::3xGFP* seedlings grown as described in (A). Sum intensity Z-projections are shown. Scale bars, 1 mm (C) Relative response of hypocotyl elongation in 3-day-old WT and *ddc* seedlings transferred to 22°C or 29°C for 3 additional days, treated with the synthetic auxin picloram (0.1 and 0.5 µM). The relative response is defined as the ratio of hypocotyl length at 29°C over 22°C of each genotype and/or treatment. (D) Quantification of hypocotyl length of WT and *yucca6-1D* seedlings, grown as described in (A). (E) Hypocotyl relative response of the seedlings shown in (D). For (C), (D), and (E), data are mean ± SEM of *n* = 3-4 independent biological replicates. Letters denote groups with significant differences from one-way analysis of variance (ANOVA) followed by Tukey’s *post hoc* tests at *P* < 0.05.

### DNA methylation is required to induce PIF4-targeted genes during thermomorphogenesis

Increases in ambient temperature induce the expression of genes involved in auxin synthesis, such as *YUCCA* (*YUC*) genes, thereby promoting hypocotyl elongation. PIF4 is a central regulator in this process, directly activating genes required for auxin synthesis like *YUC8*, *TRYPTOPHAN AMINOTRANSFERASE OF ARABIDOPSIS 1* (*TAA1*), and *CYTOCHROME P450 FAMILY 79B* (*CYP79B2*) (Franklin *et al*., 2011; Sun *et al*., 2012). Additionally, PIF4 directly induces other components of the auxin signaling pathway, such as *INDOLE-3-ACETIC ACID INDUCIBLE 19* (*IAA19*) and *IAA29* (Koini *et al*., 2009; Oh *et al*., 2012; Zhu *et al*., 2016). Since DNAme modulates auxin levels during warming (Fig. 3), we quantified the expression levels of several genes involved in auxin synthesis or signaling using reverse transcription followed by quantitative PCR (RT–qPCR). Note that we failed to detect the expected induction of these genes at warm temperatures (Supplementary Fig. S5A), as previously reported (Franklin *et al*., 2011; Sun *et al*., 2012). However, our results align with reanalyzed publicly available RNA-seq data from WT seedlings grown under similar warm temperature conditions (Supplementary Fig. S5B).

In stark contrast to decreased auxin levels observed in DNAme-deficient seedlings (Fig. 3), *YUC8*, *YUC2*, and *TAA1* transcripts were upregulated in *ddc* mutants grown at 22°C or exposed to 29°C compared to WT controls (Fig. 4A). Additionally, the transcripts of *IAA19* and *IAA29* were significantly repressed in *ddc* mutants at 22°C, but their expression remained unchanged at 29°C (Fig. 4A). Notably, we found that the *CYP79B2* transcript levels were largely downregulated in the *ddc* seedlings compared with the WT controls under both growth conditions (Fig. 4A). Since CYP79B2 catalyzes the conversion of tryptophan (Trp) into indole-3-acetaldoxime (IAOx), a precursor of the main plant auxin indole-3-acetic acid (IAA) (Zhao *et al*., 2002; Sugawara *et al*., 2009), our data implies that DNAme might regulate auxin levels by modulating the IAOx pathway.

**Fig. 4.**
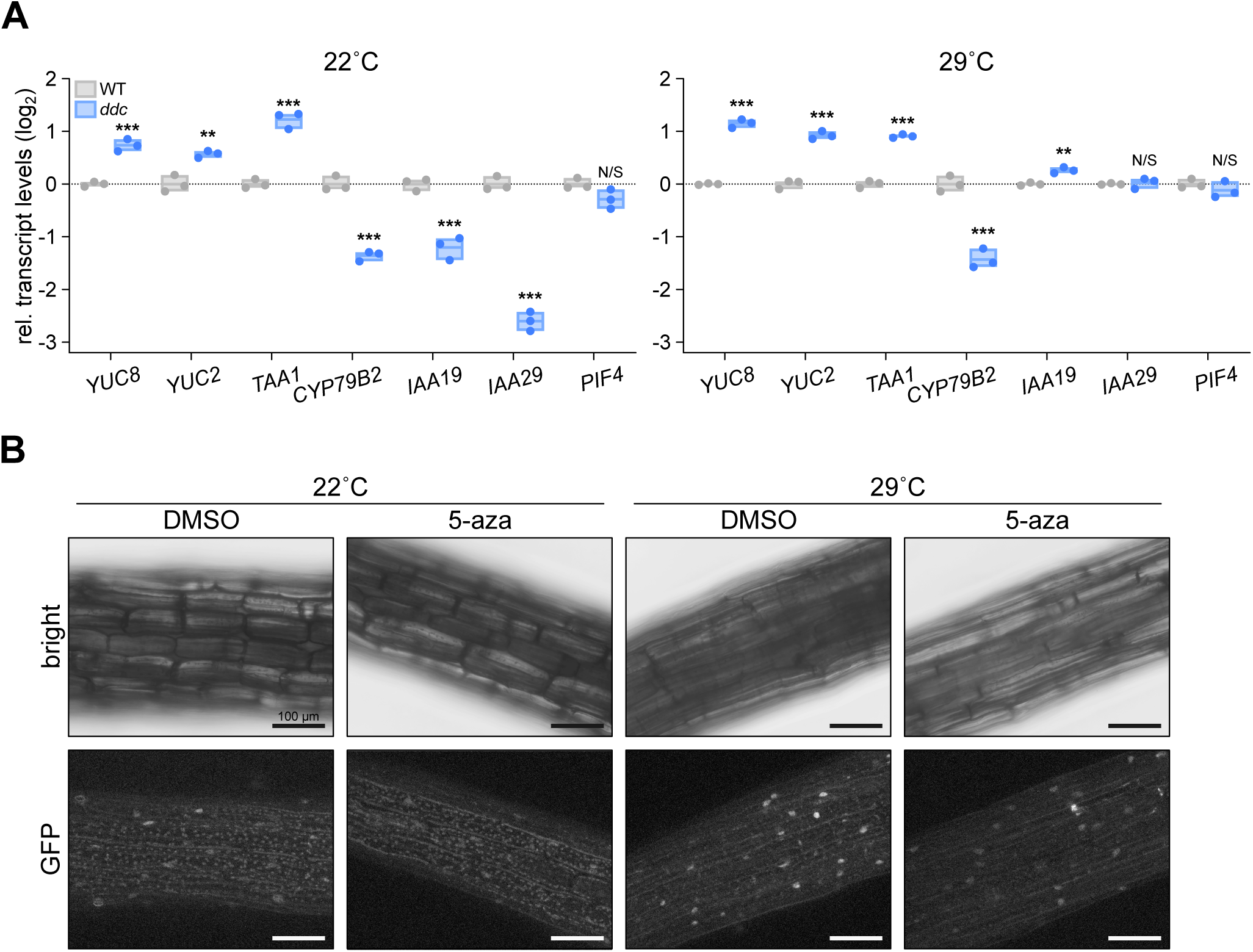
DNA methylation regulates auxin biosynthesis genes and promotes PIF4 protein levels at warm temperatures. (A) Transcript levels of *YUC8*, *YUC2*, *CYP79B2*, *TAA1*, *IAA19*, *IAA29*, and *PIF4*, quantified by RT–qPCR on 6-day-old WT and *ddc* seedlings grown at 22°C (left), or on 3-day-old WT and *ddc* seedlings transferred to 29°C for 3 additional days (right). Data are normalized to *SAND* transcript levels, and shown in log_2_ scale relative to WT (*n* = 3 independent biological replicates). The individual replicates are shown in a floating bar plot and the line depicts the mean. Statistical analysis was performed using a two-tailed Student’s *t*-test. (B) Representative pictures of live-cell imaging of *pPIF4::PIF4-GFP* seedlings transferred to 22°C or 29°C for 3 additional days, in the absence or presence of DNA methyltransferases inhibitor 5-aza (75 µM). Scale bars, 100 µm. Max intensity Z-projections are shown.

Under warm conditions, the auxin is produced within cotyledons and transported to the hypocotyl via PIN-FORMED (PIN) auxin efflux transporters (Bellstaedt *et al*., 2019), prompting us to investigate whether DNAme-deficient mutants might impact auxin distribution. As previously reported (Forgione *et al*., 2019), the *ddc* mutants negatively regulated the *PIN1*, *PIN3*, and *PIN4* transporters at 22°C (Supplementary Fig. S5C, left panel). Additionally, the auxin influx carrier *AUX1* transcript levels were downregulated under normal growth conditions (Supplementary Fig. S5D). Notably, the expression of these transporters was further repressed in *ddc* seedlings grown at 29°C (Supplementary Fig. S5C, right panel), suggesting that DNAme modulates auxin distribution during thermomorphogenesis.

Given that PIF4 directly regulates the tested genes involved in auxin biosynthesis, we investigated whether DNAme might influence *PIF4* expression. Interestingly, we found that *PIF4* transcript levels were similar in WT and *ddc* seedlings exposed to 22°C or 29°C (Fig. 4A). To explore whether DNAme might modulate PIF4 protein levels, we performed confocal microscopy on *pPIF4::PIF4-GFP* seedlings grown in the presence or absence of 5-aza at 22°C or after a shift to 29°C. As previously reported (Legris *et al*., 2017), we observed an increase in the nuclear abundance of PIF4 protein in seedlings exposed to 29°C compared to those grown at 22°C (Fig. 4B). However, in seedlings treated with 5-aza, PIF4 abundance decreased in response to 29°C (Fig. 4B; Supplementary Fig. 5E), indicating that DNAme positively regulates PIF4 protein levels. From these data, we conclude that DNAme might regulate IAOx-dependent auxin biosynthesis during thermomorphogenesis by modulating PIF4 abundance.

### DNA methylation promotes hypocotyl elongation at warm temperatures through inhibition of *SDC* expression

Plants lacking non-CG DNAme exhibit curled leaves and reduced growth, which arise from the promoter demethylation and subsequent upregulation of *SUPPRESSOR OF drm1 drm2 cmt3* (*SDC*) (Henderson and Jacobsen, 2008). To investigate whether *SDC* overexpression might be responsible for the absence of thermomorphogenic response in the *ddc* mutant, we examined hypocotyl elongation of seedlings expressing *SDC* controlled by the cassava vein mosaic virus (*CsVMV*) promoter (Tian *et al*., 2021). Using RT-qPCR, we detected similar *SDC* transcript levels between *ddc* mutants and seedlings overexpressing *SDC* (*SDCox*) (Fig. 5A). Like DNAme mutants, *SDCox* seedlings exhibited shorter hypocotyls when grown at 29°C compared to WT controls due to impaired cell elongation (Fig. 5B-D). Moreover, adding picloram partially restored the elongation defect at 29°C in *SDCox* seedlings (Supplementary Fig. S6A,B). Next, we analyzed selected targets by RT–qPCR to test whether *SDC* overexpression contributes to the deregulation of auxin-related genes observed in the DNAme-deficient seedlings. Unlike *ddc* mutants, *SDC* overexpression did not impact *YUC8* transcript levels compared to WT seedlings exposed to 29°C (Fig. 5E). However, elevated levels of *SDC* repressed *CYP79B2* and *PIN1* expression (Fig. 5E). These findings suggest that SDC negatively modulates IAOx-dependent auxin biosynthesis and transport during the warm-induced hypocotyl elongation.

**Fig. 5.**
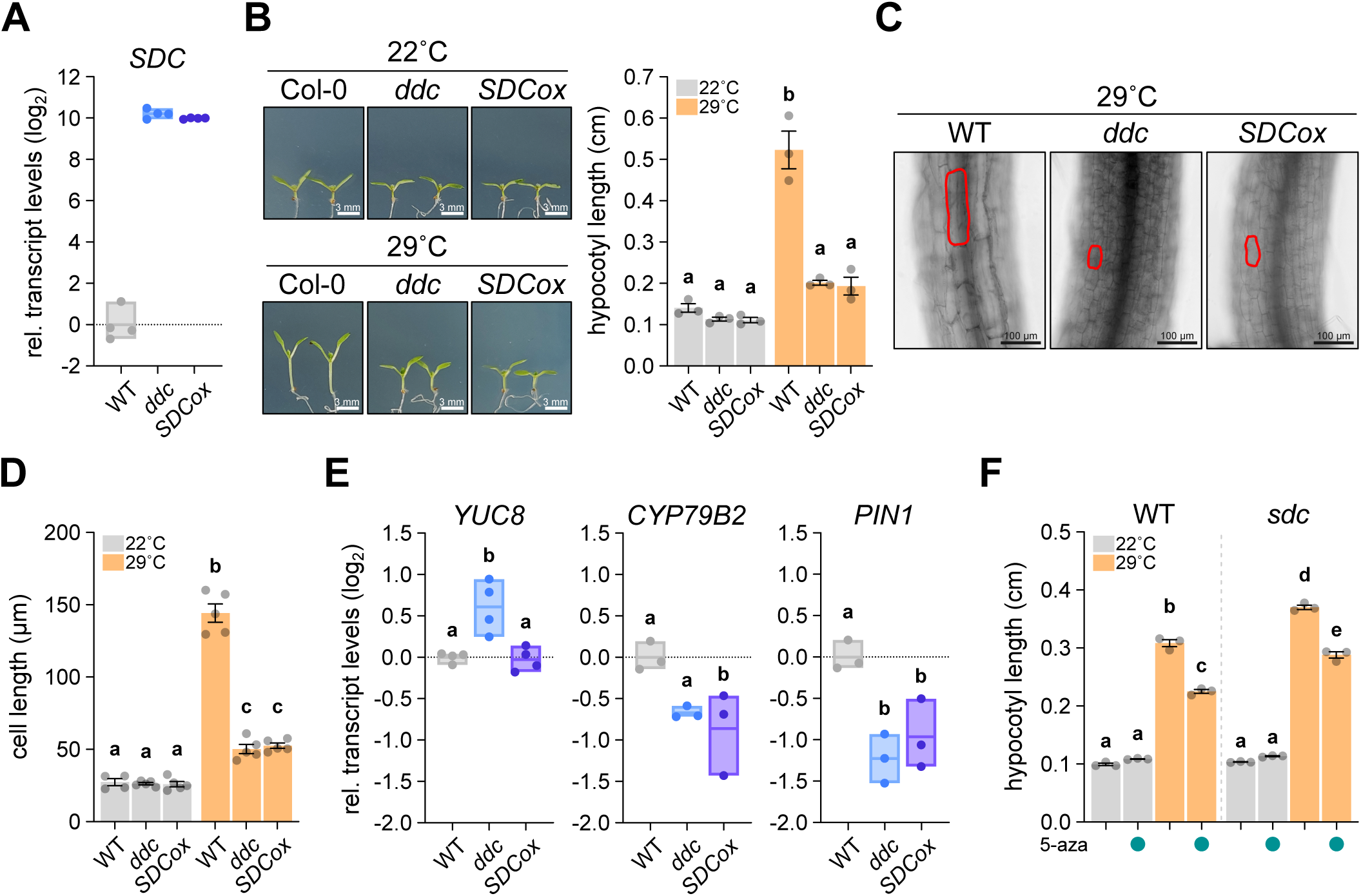
*SDC* overexpression represses thermomorphogenesis in *Arabidopsis thaliana*. (A) Transcript levels of *SDC* quantified by RT–qPCR on 6-day-old WT, *ddc*, and *SDCox* seedlings grown at 22°C. Data are normalized to *SAND* transcript levels, and shown in log_2_ scale relative to WT (*n* = 3 independent biological replicates). The individual replicates are shown in a floating bar plot and the line depicts the mean. (B) Illustrative photographs (left) and quantification of hypocotyl length (right) of 3-day-old WT, *ddc*, and *SDCox* seedlings transferred to 22°C or 29°C for 3 additional days. Scale bars, 2 mm. (C-D) Representative images (C) and quantification (D) of hypocotyl cell length from WT, *ddc*, and *SDCox* seedlings, grown as described in (B). Scale bars, 100 µm. (E) Transcript levels of *YUC8*, *CYP79B2*, and *PIN1*, quantified by RT–qPCR on 3-day-old WT, *ddc*, and *SDCox* seedlings transferred to 29°C for 3 additional days. Data are normalized to *PP2A.A3* transcript levels, and shown in log_2_ scale relative to WT (*n* = 3-4 independent biological replicates). The individual replicates are shown in a floating bar plot and the line depicts the mean. (F) Hypocotyl length measurements of 3-day-old WT and *sdc* mutant seedlings transferred to 22°C or 29°C for 3 additional days, in the absence or presence of DNA methyltransferases inhibitor 5-aza (75 µM). For (B), (D), and (F), data are mean ± SEM of *n* = 3-5 independent biological replicates. For (B), (D), (E), and (F), letters denote groups with significant differences from one-way analysis of variance (ANOVA) followed by Tukey’s *post hoc* tests at *P* < 0.05.

The role of SDC during thermomorphogenesis prompted us to test whether changes in ambient temperature regulate its expression. Using RT-qPCR, we found no significant differences in *SDC* expression when WT seedlings were grown at 22°C or 29°C (Supplementary Fig. S7A). Similar results were obtained when we reanalyzed publicly available RNA-seq of WT seedlings grown at warm temperatures (Supplementary Fig. S7B). Although *SDC* exhibits rhythmic expression specifically in *ddc2c3* mutants (Tian *et al*., 2021), *SDC* transcript levels remained unaltered in *ddc* seedlings exposed to 29°C (Supplementary Fig. S7A). These findings suggest that warm temperature does not influence *SDC* transcript levels.

Since *SDC* overexpression impairs the hypocotyl response during thermomorphogenesis, we then examined the hypocotyl length in seedlings lacking *SDC*, grown under warm temperatures in the presence or absence of 5-aza. Supporting the negative role of SDC in hypocotyl growth, we observed that *sdc* mutants showed slightly longer hypocotyls at 29°C than WT controls (Fig. 5F). However, the treatment with 5-aza resulted in similarly reduced elongation in WT and *sdc* mutants under warm conditions, compared to their untreated controls (Fig. 5F). These results suggest that the repression of *SDC* expression by DNAme plays a significant role in the temperature-induced hypocotyl elongation, but also indicates that DNAme might regulate SDC-independent processes to promote thermomorphogenesis.

### SDC inhibits hypocotyl growth during thermomorphogenesis independently of the circadian clock

To synchronize internal processes with environmental cues, the circadian clock depends on input signals that initiate transcriptional–translational feedback loops. For instance, the morning-expressed genes *CIRCADIAN CLOCK ASSOCIATED 1* (*CCA1*) and *LATE ELONGATED HYPOCOTYL* (*LHY*) repress daytime genes like *PSEUDO-RESPONSE REGULATOR 5/7/9* (*PRR5/7/9*) and the evening gene *TIMING OF CAB EXPRESSION1* (*TOC1*), which in turn suppress *CCA1* and *LHY* expression, creating a tightly regulated feedback mechanism (Huang and Nusinow, 2016). SDC affects the circadian clock by binding and destabilizing the F-box protein ZEITLUPE (ZTL), which degrades TOC1 and PRR5 in darkness (Más *et al*., 2003; Kiba *et al*., 2007). Since activated SDC inhibits TOC1 degradation (Tian *et al*., 2021), we explored whether *SDC* overexpression represses hypocotyl elongation at warm temperatures by affecting the circadian clock. To assess this, we measured the transcript levels of *CCA1* in WT, *ddc,* and *SDCox* seedlings grown at 22°C or 29°C. *CCA1* expression was repressed in *ddc* and *SDCox* seedlings compared to WT controls at ZT0 when grown at 22°C (Supplementary Fig. S8A, left panel), consistent with the notion that increased TOC1 levels inhibit *CCA1* (Gendron *et al*., 2012; Huang *et al*., 2012). However, no significant differences in *CCA1* expression were observed at 29°C between all analyzed seedlings (Supplementary Fig. S8A, right panel), suggesting the role of SDC in thermomorphogenesis might not involve circadian regulation. Supporting this notion, the treatment with 5-aza resulted in a similar inhibition of hypocotyl growth in WT and *cca1 lhy* double mutants under warm conditions, compared to their untreated controls (Supplementary Fig. S8B). Our findings suggest that SDC inhibits hypocotyl growth during thermomorphogenesis independently of the circadian rhythm regulation.

### DNA methylation regulates thermomorphogenesis partially through the modulation of gibberellin homeostasis

Gibberellins (GA) positively regulate PIF4 protein levels to promote hypocotyl elongation (Li *et al*., 2016). Since DNAme is necessary to maintain PIF4 stability at 29°C (Fig. 4), we postulated that non-CG DNAme might enhance GA content and/or activity to promote hypocotyl growth during thermomorphogenesis. To test whether mutants lacking DNAme exhibit GA deficiency, we examined the effect of GA treatment on hypocotyl elongation in *ddc* and *SDCox* seedlings grown at 29°C. Treatment with exogenous GA rescued the impaired hypocotyl growth of the *ddc* mutants at warm temperatures (Fig. 6A). Notably, *SDCox* seedlings exhibited a similar behavior compared to *ddc* mutants (Fig. 6A), suggesting that DNAme-mediated inhibition of *SDC* expression might positively modulate GA homeostasis during thermomorphogenesis. In agreement, the effect of the GA synthesis inhibitor paclobutrazol (PAC) was less pronounced in *ddc* and *SDCox* seedlings than in WT controls (Fig. 6B). Furthermore, the addition of GA restored the hypocotyl elongation defect of *dcl234* mutants during warming (Supplementary Fig. S9A). To investigate whether GA deficiency in the absence of DNAme is linked to altered GA metabolism, we next quantified the transcript levels of the GA biosynthesis genes *GA_3_OX1*, *GA_3_OX2*, and *GA_20_OX1* using RT-qPCR. Notably, we found that the expression of these selected genes was significantly repressed in *ddc* and *SDCox* seedlings exposed to 29°C (Fig. 6C). These data suggest that DNAme promotes GA biosynthesis gene expression by repressing *SDC* transcripts, thereby contributing to warm-induced hypocotyl growth.

**Fig. 6.**
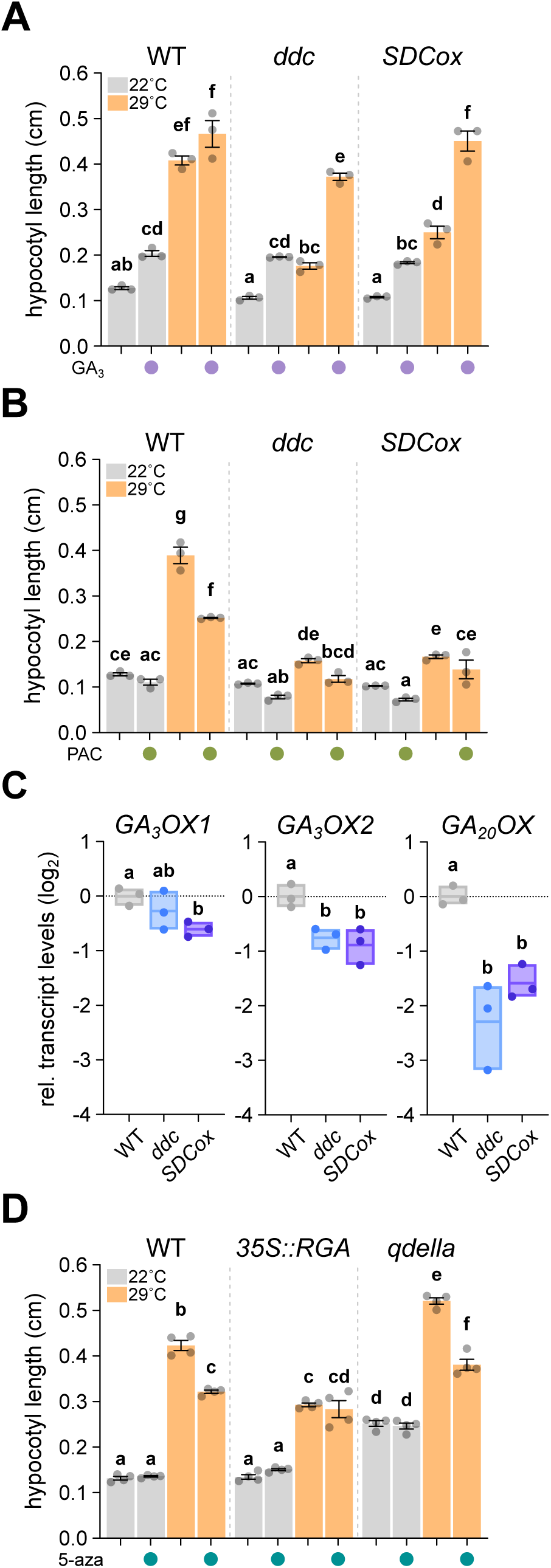
DNA methylation modulates GA metabolism to promote hypocotyl elongation during thermomorphogenesis. (A,B) Quantification of hypocotyl length of 3-day-old WT, *ddc*, and *SDCox* seedlings transferred to 22°C or 29°C for 3 days, in the presence or absence of gibberellin (GA_3_, A) or the gibberellin synthesis inhibitor paclobutrazol (PAC, B). (C) Transcript levels of *GA3OX1*, *GA3OX2*, and *GA20OX1* quantified by RT–qPCR on 3-day-old WT, *ddc*, and *SDCox* seedlings transferred to 29°C for 3 additional days. Data are normalized to *PP2A.A3* transcript levels, and shown in log_2_ scale relative to WT. The individual replicates are shown in a floating bar plot and the line depicts the mean. (D) Hypocotyl length measurements of 3-day-old WT, *35S::TAP-RGA*, and *qdella* seedlings transferred to 22°C or 29°C for 3 additional days, in the absence or presence of DNA methyltransferases inhibitor 5-aza (75 µM). For (A) to (D), data are mean ± SEM of *n* = 3-4 independent biological replicates. Letters denote groups with significant differences from one-way analysis of variance (ANOVA) followed by Tukey’s *post hoc* tests at *P* < 0.05.

GAs promote hypocotyl growth by controlling the levels of DELLA proteins, including GIBBERELLIC ACID INSENSITIVE (GAI), REPRESSOR OF ga1–3 (RGA), RGA-like 1 (RGL1), RGL2, and RGL3 (Shani *et al*., 2024). To further explore the role of DNAme on GA metabolism, we measured hypocotyl elongation in seedlings overexpressing *DELLA* (*35S::TAP-RGA* and *35S::TAP-GAI*), and the quintuple *della* mutant (*qdella*) grown in the presence or absence of 5-aza at either 22°C or 29°C. Although seedlings overexpressing *RGA* or *GAI* exhibited short hypocotyls at 29°C, they were insensitive to the presence of 5-aza, unlike the WT controls (Fig. 6D; Supplementary Fig. S9B). Consistent with the *sdc* mutants (Fig. 5F), 5-aza treatment resulted in similarly impaired hypocotyl growth in WT and *qdella* seedlings when grown at 29°C, compared to their untreated controls (Fig. 6D). Taken together, our findings suggest that non-CG DNAme promotes hypocotyl elongation through the induction of GA biosynthesis, but also imply that DNAme might modulate other molecular factors to regulate thermomorphogenesis.

## Discussion

Thermomorphogenesis encompasses a range of developmental and molecular mechanisms that enable plants to adapt to warm environmental temperatures. This process is tightly regulated by integrating light signals, phytohormones, and circadian clock pathways (Casal and Balasubramanian, 2019; Ding *et al*., 2020). Additionally, epigenetic regulatory mechanisms are crucial in modulating plant thermomorphogenic responses (Tasset *et al*., 2018; He *et al*., 2022; Kim *et al*., 2023; Zhou *et al*., 2024). Here, we have identified the conserved epigenetic mark DNA methylation as a key regulator of the plant response to warm temperatures. Indeed, the absence of DNAme impairs hypocotyl elongation during thermomorphogenesis (Figs. 1 and 2). Furthermore, we demonstrate that DNAme promotes increments in auxin levels in the hypocotyl by modulating gibberellin metabolism and auxin transporters in an SDC-dependent manner (see model in Fig. 7).

**Fig. 7.**
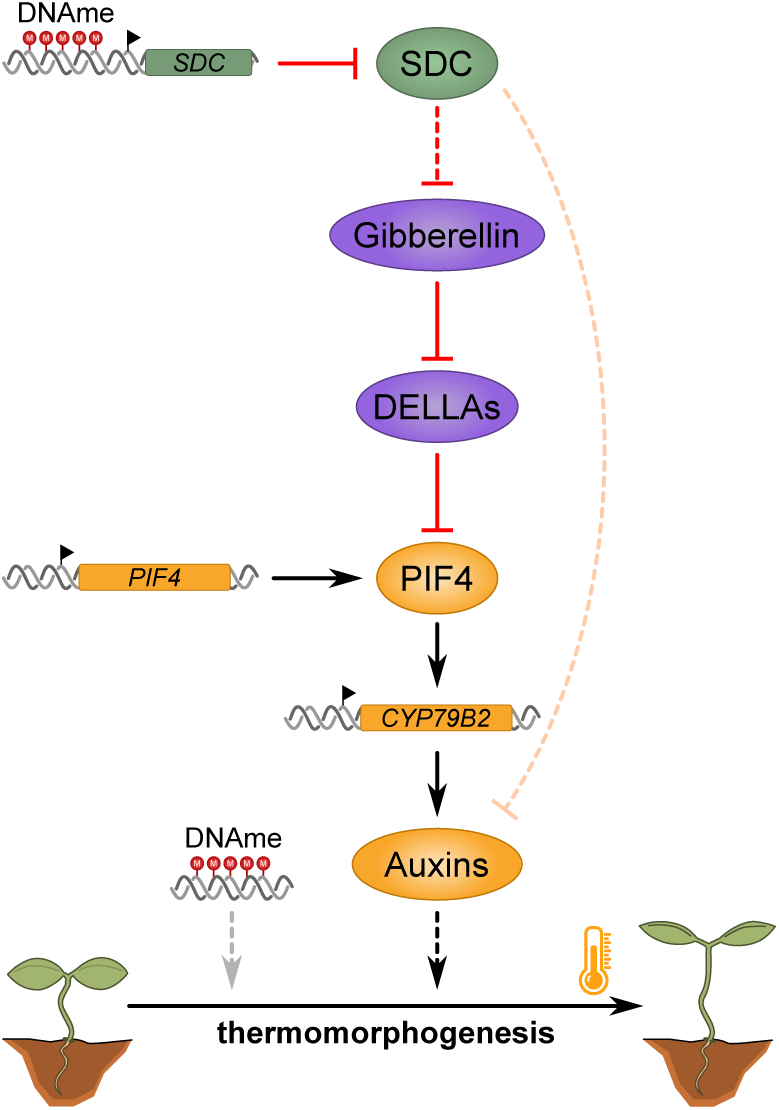
Proposed model for the role of DNAme in the regulation of hypocotyl elongation during thermomorphogenesis. DNAme represses *SDC* expression to promote GA synthesis and prevent DELLA-mediated degradation of PIF4, thereby enhancing IAOx-dependent auxin synthesis and hypocotyl growth. Additionally, *SDC* repression affects other aspects of auxin metabolism, such as auxin transport, in a PIF4-independent manner (orange dashed line). Besides *SDC* repression, DNAme might also influence additional regulatory pathways to promote temperature-induced hypocotyl growth (gray dashed line).

During thermomorphogenesis, hypocotyl growth relies on increased auxin activity driven by PIF4-mediated induction of genes involved in auxin biosynthesis (Gray *et al*., 1998; Franklin *et al*., 2011; Sun *et al*., 2012). Consistently, we found that the absence of DNAme reduces auxin activity and destabilizes PIF4 protein in the hypocotyl of seedlings exposed to warm temperatures (Figs. 3 and 4; Supplementary Fig. 5D). Notably, DNAme regulates PIF4 levels without affecting its transcription (Fig. 4), unlike other epigenetic marks previously reported (Tasset *et al*., 2018). Moreover, the exogenous application of an auxin analog partially rescues the hypocotyl growth defect of *ddc* mutants (Fig. 3C; Supplementary Fig. S4). We also found that DNA hypomethylation inhibits the expression of auxin transporters (Supplementary Fig. S5C), thereby disrupting the cotyledon-to-hypocotyl transport of IAA needed for cell elongation at warm temperatures (Bellstaedt *et al*., 2019). Interestingly, this auxin accumulation in cotyledons might trigger alternative developmental responses, such as the curled leaf phenotype seen in the *ddc* mutants (Forgione *et al*., 2019, 2022).

The biosynthesis of the main auxin IAA involves several pathways, including the conversion of Trp to IAOx by the enzymes CYP79B2 and CYP79B3 (Casanova-Sáez *et al*., 2021). We found that *CYP79B2* expression is significantly repressed in the *ddc* mutants (Fig. 4A). This suggests that DNAme not only regulates auxin transport but also influences IAA levels by modulating the IAOx pathway. Supporting this, the double *cyp79b2 cyp79b3* mutant, which exhibits normal IAA content under normal growth conditions, fails to elongate the hypocotyl during warming (Zhao *et al*., 2002). Moreover, channeling IAOx to IAA biosynthesis promotes temperature-induced hypocotyl growth (Sugawara *et al*., 2009; Kong *et al*., 2015).

Although the absence of DNAme diminished PIF4 protein levels, several direct PIF4 target genes are induced in the *ddc* mutants (Fig. 4), suggesting that DNAme might regulate their expression independently of PIF4 activity. For instance, DNAme at *YUC2* and *IAA* promoters inhibits their transcription, and its removal induces *YUC2* expression at warm temperatures (Fonouni-Farde *et al*., 2022; Forgione *et al*., 2022). DNAme also modulates the expression of components of the Polycomb repressive complex 2 (PRC2) that deposits the repressive mark H3K27me3, a required modification for *YUC8* induction during thermomorphogenesis (Forgione *et al*., 2022; Kim *et al*., 2023). This suggests an indirect role of DNAme in modulating the expression of PIF4 targets.

The developmental changes observed in non-CG DNAme-deficient mutants result from *SDC* induction (Henderson and Jacobsen, 2008; Tian *et al*., 2021). Our findings support the notion that DNAme regulates *SDC* expression to control thermomorphogenesis, as evidenced by the altered hypocotyl elongation in seedlings overexpressing *SDC* (Fig. 5B). Additionally, *SDC* overexpression negatively regulates *CYP79B2* and *PIN1* expression, similar to what is detected in *ddc* mutants (Fig. 5E). However, the observation that DNAme influences hypocotyl growth even in the absence of *SDC* suggests that other molecular factors are also modulated by DNAme to regulate thermomorphogenesis (Fig. 5F). For instance, *YUC8* transcript levels are deregulated in *ddc* mutants but remain unaltered in *SDCox* seedlings (Fig. 5E). Therefore, while SDC plays a significant role, the broader regulatory impact of DNAme on additional pathways should not be overlooked (Fig. 7).

Although SDC mediates most of the effects of DNA hypomethylation on hypocotyl elongation during thermomorphogenesis, we found that its expression is not directly influenced by warm temperatures (Supplementary Fig. 7). Nevertheless, high temperatures can release the epigenetic silencing of *SDC*, leading to its induction, particularly in young developing leaves (Sanchez and Paszkowski, 2014). This suggests that warm temperatures are not sufficient to influence the presence (or absence) of specific factors required for *SDC* expression. Beyond DNAme, *SDC* expression is repressed by the ATPase-containing MICRORCHIDIA (MORC) protein, reinforcing the silencing of DNA-methylated genes (Moissiard *et al*., 2012). Moreover, the IMITATION SWITCH (ISWI) complex regulates nucleosome occupancy over *SDC*, resulting in *SDC* induction independently of the DNAme status (Zhang *et al*., 2023). However, it remains unclear whether specific environmental stimuli might influence the DNAme pattern or the binding of MORC and/or ISWI to the *SDC* promoter to regulate its expression and confer an adaptative advantage. Supporting this notion, *SDC* expression is significantly induced in several Arabidopsis natural accessions (Kawakatsu *et al*., 2016; Tian *et al*., 2021), implying that SDC likely contributes to the regional adaptation of different ecotypes. It also raises the possibility that these accessions exhibit an altered warming response, warranting further investigation.

SDC encodes an F-box containing protein, with its only known targets being factors involved in controlling the circadian clock, such as ZTL (Tian *et al*., 2021). Notably, ZTL regulates PIF4 activity and expression to modulate plant response to warm temperatures (Seo *et al*., 2023). This raises the possibility that SDC represses hypocotyl elongation by promoting ZTL degradation. In stark contrast, we found that the impaired hypocotyl growth in DNAme-deficient mutants during thermomorphogenesis seems independent of circadian clock regulation (Supplementary Fig. S8). Furthermore, seedlings lacking DNAme do not exhibit altered *PIF4* expression (Fig. 4A), unlike *ztl* mutants (Seo *et al*., 2023).

Endogenous phytohormones integrate temperature information to regulate hypocotyl growth during thermomorphogenesis (Lu *et al*., 2021). For instance, GA levels rise in aerial tissues at warm temperatures due to enhanced root-to-shoot translocation of an inactive GA precursor, thereby promoting hypocotyl growth through a DELLA-dependent mechanism (Camut *et al*., 2019). In agreement, we found that *SDC* overexpression represses GA synthesis, and exogenous application of the phytohormone restores hypocotyl elongation in *ddc* and *SDCox* seedlings (Fig. 6). Given that DELLA interaction leads to PIF4 degradation (Li *et al*., 2016), we propose a model in which DNAme promotes GA synthesis in an SDC-dependent manner to stabilize PIF4 protein levels, contributing to hypocotyl growth (Fig. 7). In line with this model, elevated levels of DELLA inhibit temperature-induced hypocotyl elongation independently of the global DNAme pattern (Fig. 6D and Supplementary Fig. S9B). Although a direct regulation of GA biosynthesis genes by DNAme cannot be excluded, *SDC* overexpression also negatively regulates the expression of *GA_3_OX1*, *GA_3_OX2*, and *GA_20_OX1* (Fig. 6C). Given that SDC is an F-box protein, these findings suggest that DNAme might regulate GA biosynthesis through the degradation of a yet unidentified factor. One potential target is the histone H3K27 demethylase RELATIVE OF EARLY FLOWERING 6 (REF6), as REF6 is required to promote hypocotyl elongation by inducing *GA_20_OX1* expression (He *et al*., 2022). In line with this, *ref6* mutants exhibit reduced expression of *PIN* transporters (Wang *et al*., 2019), similar to the patterns observed in *SDC*-overexpressing seedlings (Fig. 6C). Other potential targets of SDC might include the TEOSINTE BRANCHED 1/CYCLOIDEA/PCF (TCP) transcription factors TCP14 and TCP15, which also regulate GA biosynthesis to promote thermomorphogenesis (Ferrero *et al*., 2019). Future research should focus on identifying the specific targets of SDC that contribute to auxin and/or GA metabolism regulation.

Overall, our results highlight the importance of epigenetic modifications, specifically DNA methylation, in regulating plant growth responses to environmental stimuli. This expands our knowledge beyond genetic factors, emphasizing how epigenetic changes can influence phenotypic plasticity.

## Supplementary data

Supplementary Fig. S1. DNA methylation promotes hypocotyl elongation during thermomorphogenesis.

Supplementary Fig. S2. DNA methylation does not influence petiole elongation during thermomorphogenesis.

Supplementary Fig. S3. The reagent 5-azacytidine inhibits hypocotyl elongation of tobacco plants exposed to elevated temperatures.

Supplementary Fig. S4. Exogenous auxin can partially rescue the hypocotyl growth of DNAme-deficient mutants during thermomorphogenesis.

Supplementary Fig. S5. DNA methylation is required to induce PIF4-regulated genes during thermomorphogenesis.

Supplementary Fig. S6. Exogenous auxin partially rescues deficient hypocotyl growth in seedlings overexpressing *SDC*.

Supplementary Fig. S7. *SDC* expression is unaffected by warm temperatures.

Supplementary Fig. S8. *SDC* overexpression represses warm-induced hypocotyl elongation independently of the circadian clock.

Supplementary Fig. S9. DELLA proteins mediate DNA methylation-dependent hypocotyl growth during thermomorphogenesis.

Supplementary Table S1: Set of primers used in this study for RT-qPCR experiments.

## Acknowledgments

The authors are grateful to the plant community for their willingness to share seed material with us. The authors thank Camila Seimandi and Dr. Carlos M. Figueroa for their help with the setting of some experiments. We thank Dr. Raquel L. Chan for reviewing an earlier draft of this paper.

## Author contributions

MC: conceptualization; LB and MC: design; MG, EG, GJV, AL, and MC: performing experiments; LB and MC: funding acquisition; MC: visualization; MC: writing-review and editing. All the authors contributed to the data analysis and discussion, and reviewed and revised the manuscript.

## Data availability

All data supporting the findings of this study, including supplementary materials, are available from the corresponding author upon request.

## Supplementary figure legends

**Supplementary Fig. S1. DNA methylation promotes hypocotyl elongation during thermomorphogenesis.**

(A) Top left, scheme of the temperature conditions used to evaluate hypocotyl length. Bottom left, illustrative photographs of WT, *ddc*, and *pif4-2* hypocotyls grown at 22°C or 29°C. Scale bars, 2 mm. Right, quantification of hypocotyl length of WT, *ddc*, and *pif4-2* seedlings grown at 22°C or 29°C for 5 days. B-E) Hypocotyl length in 3-day-old WT (Col-0), and the indicated mutants in DNA methylation maintenance (B and C), small RNA generation (D), or DNA demethylation (E) seedlings transferred 22°C or 29°C for 3 additional days. For (B) to (E), data are mean ± SEM of *n* = 3-4 independent biological replicates. Letters denote groups with significant differences from one-way analysis of variance (ANOVA) followed by Tukey’s *post hoc* tests at *P* < 0.05. *ddc*, *drm1-2 drm2-2 cmt3-11*; *ddc2c3*, *drm1-2 drm2-2 cmt2-3 cmt3-11*; *dds4*, *drm1-2 drm2-2 kyp-6*; *dcl234*, *dcl2-1 dcl3-1 dcl4-2t*; *rdd-1*, *ros1-3 dml2-1 dml3-1*; *rdd-2*, *ros1-4 dml2-2 dml3-2*; *drdd*, *dme ros1-4 dml2-2 dml3-2 dme^DD7pro^*.

**Supplementary Fig. S2. DNA methylation does not influence petiole elongation during thermomorphogenesis.**

(A) Top, schematic representation of the growth conditions used to analyze petiole length. Bottom, representative images of 9-day-old WT, *ddc*, and *pif4-2* plants transferred to 22°C or 29°C for 7 d. Scale bars, 5 mm. (B) Quantification of petiole length from plants shown in (A). (C) Petiole length of WT, *ddc*, and *pif4-2* plants at 29°C relative to 22°C. Data are mean ± SEM from *n* independent biological replicates. Letters denote groups with significant differences from one-way analysis of variance (ANOVA) followed by Tukey’s *post hoc* tests at *P* < 0.01.

**Supplementary Fig. S3. The reagent 5-azacytidine inhibits hypocotyl elongation of tobacco plants exposed to elevated temperatures.**

(A) Top, scheme of the temperature conditions used to evaluate hypocotyl length in tobacco plants. Bottom, illustrative photographs of 7-day-old *N. benthamiana* plants transferred to 22°C or 29°C for 4 additional days, in the absence or presence of 5-azacytidine 75 µM (5-aza). Scale bars, 3 mm. (B) Quantification of hypocotyl length in WT seedlings grown as described in (A). Data are mean ± SEM of *n* = 6 independent biological replicates. Letters denote groups with significant differences from one-way analysis of variance (ANOVA) followed by Tukey’s *post hoc* tests at *P* < 0.05.

**Supplementary Fig. S4. Exogenous auxin can partially rescue the hypocotyl growth of DNAme-deficient mutants during thermomorphogenesis.**

Quantification of hypocotyl length in 3-day-old WT, *ddc*, and *pif4-2* seedlings transferred to 22°C or 29°C for 3 additional days, in the absence or presence of the synthetic auxin picloram (0.1 and 0.5 µM). Data are mean ± SEM of *n* = 3 independent biological replicates. Letters denote groups with significant differences from one-way analysis of variance (ANOVA) followed by Tukey’s *post hoc* tests at *P* < 0.05.

**Supplementary Fig. S5. DNA methylation is required to induce PIF4-regulated genes during thermomorphogenesis.**

(A) Transcript levels of *YUC8*, *YUC2*, *CYP79B2*, *TAA1*, *IAA19*, *IAA29*, and *PIF4*, quantified by RT–qPCR on 3-day-old WT transferred to 22°C or 29°C for 3 additional days. (B) Table showing the expression values of multiple auxin-related genes, as shown in Fig. 4. Significant differences are highlighted in bold. Data were retrieved from RNA-seq analyses and are shown as log_2_ fold change of 28°C over 22°C. (C) Transcript levels of *PIN1*, *PIN3*, *PIN4*, and *PIN7*, quantified by RT–qPCR on 6-day-old WT and *ddc* seedlings grown at 22°C (left), or on 3-day-old WT and *ddc* seedlings transferred to 29°C for 3 days (right). (D) Transcript levels of *AUX1* quantified by RT–qPCR on WT and *ddc* seedlings grown as described in (C). (E) Quantification of nuclei intensity from live-cell imaging of *pPIF4::PIF4-GFP* seedlings transferred to 22°C or 29°C for 3 additional days, in the absence or presence of DNA methyltransferases inhibitor 5-aza (75 µM), as in Fig. 4B. Data are mean ± SEM of *n* = 4 independent biological replicates. For (A), (C), and (D), data are normalized to *SAND* (A and C) or *PP2A.A3* (D) transcript levels, and shown in log_2_ scale relative to WT (*n* = 3 independent biological replicates). The individual replicates are shown in a floating bar plot, and the line depicts the mean. For (A), (C), (D), and (E), statistical analysis was performed using a two-tailed Student’s *t*-test. a.u., arbitrary units.

**Supplementary Fig. S6. Exogenous auxin partially rescues deficient hypocotyl growth in seedlings overexpressing *SDC*.**

(A) Quantification of hypocotyl length in 3-day-old WT, *ddc*, and *SDCox* seedlings transferred to 22°C or 29°C for 3 additional days, in the absence or presence of the synthetic auxin picloram (0.1 and 0.5 µM). (B) Relative response of hypocotyl elongation of the seedlings shown in (A). The relative response is defined as the ratio of hypocotyl length at 29°C over 22°C of each genotype and/or treatment. For (A) and (B), data are mean ± SEM of *n* = 3 independent biological replicates. Letters denote groups with significant differences from one-way analysis of variance (ANOVA) followed by Tukey’s *post hoc* tests at *P* < 0.05.

**Supplementary Fig. S7. *SDC* expression is unaffected by warm temperatures.**

(A) Transcript levels of *SDC* quantified by RT–qPCR on 3-day-old WT and *ddc* seedlings transferred to 22°C or 29°C for 3 additional days. Data are normalized to *PP2A.A3* transcript levels, and shown in log_2_ scale relative to WT at 22°C (*n* = 4 independent biological replicates). The individual replicates are shown in a floating bar plot and the line depicts the mean. Letters denote groups with significant differences from one-way analysis of variance (ANOVA) followed by Tukey’s *post hoc* tests at *P* < 0.05. (B) Coverage plots showing transcripts in 3-day-old WT seedlings transferred to 22°C or 28°C for 3 additional days, from two independent biological replicates. Reads are presented as counts per million (CPM). Genomic coordinates are shown in base pairs (bp).

**Supplementary Fig. S8. *SDC* overexpression represses warm-induced hypocotyl elongation independently of the circadian clock.**

(A) Transcript levels of *CCA1* quantified by RT–qPCR on 6-day-old WT, *ddc*, and *SDCox* seedlings grown at 22°C (left), or on 3-day-old WT, *ddc*, and *SDCox* seedlings transferred to 29°C for 3 additional days (right). Data are normalized to *PP2A.A3* transcript levels, and shown in log_2_ scale relative to WT. The individual replicates are shown in a floating bar plot and the line depicts the mean. (B) Hypocotyl length measurements of 3-day-old WT and *cca1 lhy* mutant seedlings transferred to 22°C or 29°C for 3 additional days, in the absence or presence of DNA methyltransferases inhibitor 5-aza (75 µM). For (A) and (B), data are mean ± SEM of *n* = 3-4 independent biological replicates. Letters denote groups with significant differences from one-way analysis of variance (ANOVA) followed by Tukey’s *post hoc* tests at *P* < 0.05.

**Supplementary Fig. S9. DELLA proteins mediate DNA methylation-dependent hypocotyl growth during thermomorphogenesis.**

(A) Quantification of hypocotyl length of 3-day-old WT and *dcl234* seedlings transferred to 22°C or 29°C for 3 days, in the presence or absence of gibberellin (GA_3_). (B) Hypocotyl length measurements of 3-day-old WT and *35S::TAP-GAI* seedlings transferred to 22°C or 29°C for 3 additional days, in the absence or presence of DNA methyltransferases inhibitor 5-aza (75 µM). For (A) and (B), data are mean ± SEM of *n* = 3 independent biological replicates. Letters denote groups with significant differences from one-way analysis of variance (ANOVA) followed by Tukey’s *post hoc* tests at *P* < 0.05.

